# Thymic involution as an adaptive schedule for combating diverse pathogens

**DOI:** 10.1101/2024.12.21.629868

**Authors:** Yoh Iwasa, Kosei Matsuo

## Abstract

Diverse naive T cells, produced in the thymus, migrate to peripheral tissues and help suppress pathogens upon activation. As we age, the thymus shrinks, reducing both the supply and diversity of naive T cells, which increases the time required to establish immunity against new pathogens. This process, called thymic involution, is considered a form of aging. In this study, we explore whether thymic involution might be an adaptive strategy. Over time, the immune system accumulates memory cells, reducing the need for new naive T cells to combat unfamiliar pathogens. We examine how the optimal rate of naive T cell production, *h*(*t*), declines with age, taking into account the cost of maintaining thymic activity. When pathogen encounter rates are equal, *h*(*t*) decreases exponentially with age. As pathogen encounter rates rise, the initial value *h*(0) increases, but the rate declines more rapidly with age, reducing overall production. A higher pathogen diversity, a greater cost of fighting new pathogens, and a lower cost of maintenance increase *h*(*t*). When pathogen encounter rates vary, *h*(*t*) decreases with age according to a power function.

**Highlights:**

1. The thymus generates diverse naive T cells to recognize a wide range of pathogens.
2. Thymic involution is considered an adaptive strategy, factoring in maintenance cost.
3. If pathogen encounters are frequent, the rate starts high but quickly declines.
4. The rate increases with the pathogen risk and decreases with the thymic cost.
5. If pathogens vary in abundances, the rate decreases as a power function of age.

## 1. Introduction

The thymus produces immune cells that are key agents of adaptive immunity (Parham 2021). To ensure that adaptive immunity can recognize a wide variety of pathogens, the thymus randomly generates a vast number of cells, each capable of recognizing a specific antigen, while eliminating those that react to the host’s own substances. Naive T cells produced in the thymus migrate to peripheral tissues, where they become activated by foreign antigens and help stimulate B cells, cytotoxic T cells, neutrophils, and other immune cells to suppress pathogenic substances. After these adaptive immune responses successfully suppress the pathogens, memory cells are formed and remain in the body, ready to combat the same antigens if they reinfect in the future.

The production of naive T cells in the thymus decreases with age. This is closely related to the decline of the function of the thymus, known as “thymic involution” (Lewkiewicz et al. 2019). According to a quantitative modelling taking into account many processes in the thymus, decrease in the thymic cell number starts early and by the age of 18, the number of cells decreases to about one-tenth of the level found in an infant (Kulesh et al. 2024). The exponential decrease in thymic output starts at birth, but the decrease slows down in older ages (Mitchell et al. 2010). Situation is similar in nonhuman animals, such as rats or dogs, but the decline of thymic function occurs faster than human (Fujiwara et al. 2012; Appay and Sauce 2014).

As the body volume increases, the total number of naive T cells observed in the peripheral pool increases, probably due to proliferation of naive T cells (Bains et al. 2009; Gorony et al. 2015). They may stay quiescent in a partially differentiated state in periphery (Gorony et al. 2015), although recent and detailed model for naive T cell dynamics concluded that homeostatic regulation of naive T cell number in periphery (such as density-dependent survivorship) does not operate for naive T cells in periphery (Rane et al. 2022), many being in spleen and lymph nodes (Ganusov and de Boer 2007). However, cell proliferation cannot increase the diversity of naive T cells with respect to their recognition ability of the adaptive immunity. The decline in the thymic function causes age-dependent decrease in the diversity of thymic output (Collins et al. 2024). Consequently, the time required to establish immunity against new pathogens increases.

Thymic involution has been discussed as an example of immuno-senescence, which refers to the gradual aging of the immune system (Sidler et al. 2013; Palmer et al. 2018). However, since the decrease in thymic size and the supply rate of naive T cells from the thymus to periphery starts before reproductive ages, we believe that it is likely an adaptive strategy of the immune system, rather than a result of the degradation of biological functions with age (i.e., ageing; Hamilton 1966; Sun et al. 2023).

In this study, we explore the possibility that thymic involution is an adaptive strategy for the immune system using simple mathematical models.

As an individual ages and encounters various pathogens, the immune system copes with them, accumulating memory cells for these antigens. Consequently, the need to cope with novel antigens, which may be potential pathogens, decreases as the individual grows older. On the other hand, the thymus must continue generating some naive T cells to respond to antigens that have not yet been encountered, including pathogens that have mutated from previously encountered strains. We examine how the optimal rate of naive T cell production, *h*(*t*), decreases with age *t*, taking into account the costs associated with maintaining the thymus’s capacity to produce a high number of naive T cells and the risk of encountering previously unknown pathogens. The scheme of the model is illustrated in Fig. 1.

**Figure 1.**
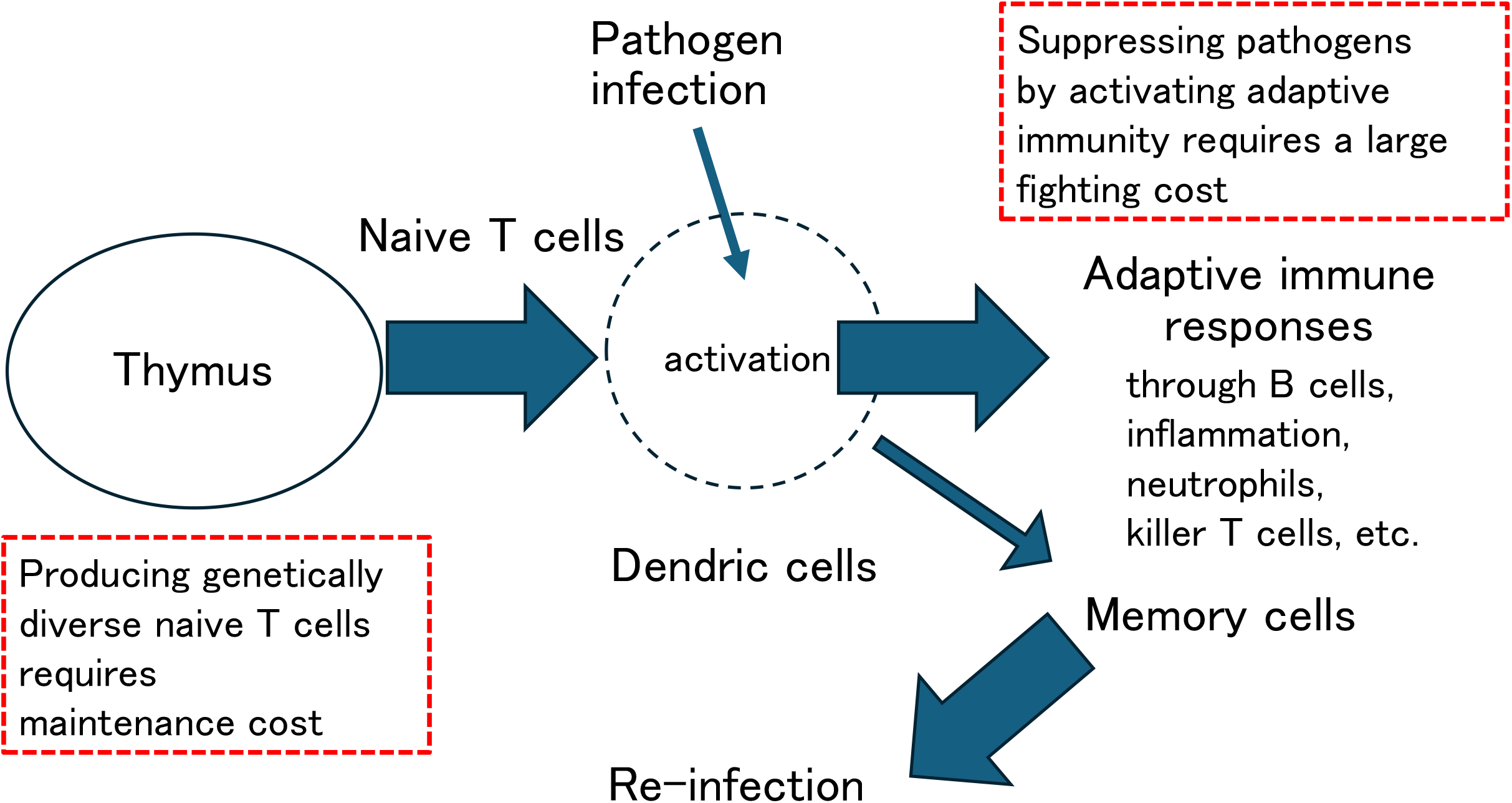
Scheme of the model. Refer to the main text for explanations.

The cost of fighting a pathogen infection would be greater if adaptive immune responses are delayed due to reduced supply of naive T cells from the thymus, allowing pathogens to proliferate. Therefore, maintaining a large thymus and a rapid supply rate of naive T cell production helps reduce the “fighting cost.” On the other hand, maintaining a large thymus and a high rate of naive T cell production comes with a cost (maintenance cost). The balance between fighting costs and maintenance costs may determine the optimal level of naive T cell production, which decreases as one ages and accumulates more memory cells in the body.

The analysis shows that the optimal rate of naive T cell production *h*(*t*) decreases with age, either as an exponential function or as a power function, depending on the variability in the pathogen encounter rate. We discuss how the rate *h*(*t*) changes in relation to the pathogen encounter rate in the environment, the risks associated with fighting with pathogens, and the costs of maintaining thymic activity.

## 2. Model

Suppose there exist many different antigens that are potentially pathogenic, which we denote *i* = 1,2,3, …., *n*, where *n* is the total number of antigens. The host encounters these antigens at random. Some are pathogenic and can invade the body and proliferate. However, others are not pathogenic, either because they cannot proliferate in the host body or because they are obstructed by the barrier functions of the skin and other epithelial tissues, or by the mechanisms of innate immunity. When the host encounters pathogen that could not be suppressed by the innate immunity, it needs the actions of adaptive immunity to prevent the harm. Let *f*_*i*_ be the rate of encounter with antigens of type *i* that requires the action of adaptive immunity (*i* = 1,2, …, *n*).

Adaptive immunity is a mechanism by which immune cells specific to a given antigen proliferate to fight against the disease. After the infection is resolved, the host retains memory cells that can be activated if the same pathogen reinfects the host in the future. When a host encounters a pathogen and retains memory cells reactive to it, the host can respond to reinfection relatively quickly. However, if the host is infected by newly invaded pathogens, the adaptive immunity must be activated. The probability for the antigen to be the one not encountered before is 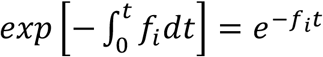, where *t* is the age of the host.

The process of fighting with a novel pathogen begins with the activation of specific naive T cells by dendritic cells in the periphery, followed by the activation of various immune cells, including B cells and cytotoxic T cells, which together trigger immune responses. We assume that the novel pathogen will eventually be suppressed, leading to the formation of memory cells capable of responding to future reinfections.

If the supply rate of naive T cells from the thymus is low, the time taken for specific naive T cells to recognize the pathogen would be delayed, allowing the pathogens to proliferate and increasing the cost of suppressing them. To represent this idea, we assume that the cost for fighting with a novel pathogen is inversely proportional to the rate of naive T cell supply: 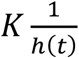, where the proportionality coefficient *K* indicates the magnitude of harm caused by the infection of a novel pathogen.

Combining these arguments, the amount of cost the host must pay in suppressing the pathogens that encounter within a short time interval of Δ*t* is the cost of fighting with a single novel pathogen 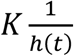 multiplied by the probability of encountering a novel pathogen that requires the activation of adaptive immunity: 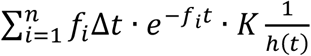, which indicates “fighting cost.” This decreases with supply rate of naive T cells *h*(*t*).

We also consider the cost for maintaining thymic activity. In the thymus, the training of immune cells takes place. The cells that are to recognize a large diversity of antigen-recognizing sites undergo hypermutation, a specific process that induces mutations in certain portions of genes at an exceptionally high rate. This process combines recombination and gene conversion, resulting in cells with a large variety of genes in the regions coding for the specificity of antigen recognition. Following this, a “negative selection” procedure occurs, during which cells that recognize the body’s own proteins and other materials are eliminated. The processes of proliferation with a high mutation rate and negative selection occur alternately over multiple cycles, ultimately producing naive T cells with a large diversity in their specificity of recognition. Hence, maintaining the thymus, which continues to produce naive T cells, involves a significant biological cost.

The maintenance cost increases as the supply rate of naive T cells *h*(*t*) increases. For simplicity, we assume that it is directly proportional to *h*(*t*) with a proportionality constant of *m*. Therefore, the cost of maintaining the rate of naive T cell supply is *mh*(*t*)Δ*t*.

Combining these two, the total cost associated with the defense action per unit time is as follows:

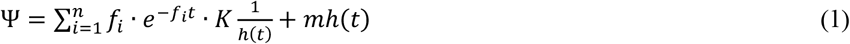

The first and second terms on the right-hand side represent the fighting cost and maintenance cost per unit time, respectively. The optimal value of *h*(*t*) is the one that minimizes Ψ, which can be calculated from setting the derivative equal to zero: 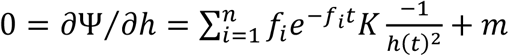, which leads to

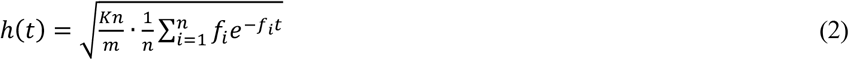

We can interpret *K* and *m* as the costs of fighting and maintenance, respectively. *n* is the diversity of antigens that are potentially pathogenic. The term 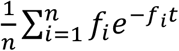 is the quantity 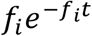 averaged over multiple antigens. The last quantity is a decreasing function of age *t*. Consequently, the optimal rate of naive T cell supply from the thymus, as described in Eq. (2), decreases with age. This decline may provide an explanation for thymic involution. In the following sections, we examine the dependence on parameters and the age-related changes in detail.

## 3. Optimal rate of producing naive T cells decreases with age

In this and next sections, we discuss the parameter dependence of the optimal rate of naive T cell production, as given in Eq. (2) of the last section. We start to consider the case in which different antigens have the same *f*_*i*_, the rate of encounter with pathogens in the environment that require action of adaptive immunity. We denote 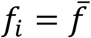, for *i* = 1,2, …, *n*. The formula for the optimal *h*(*t*) given by Eq. (2) becomes as follows:

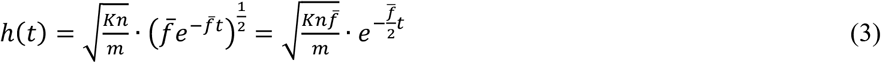

This implies that the optimal rate of naive T cell supply rate decreases with age *t* following an exponential function of age *t*.

### 3.1 Rate of encountering pathogens in the environment

Fig. 2A illustrates how the optimal rate of naive T cells declines with the age for several different values of encounter rate 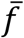.

**Figure 2.**
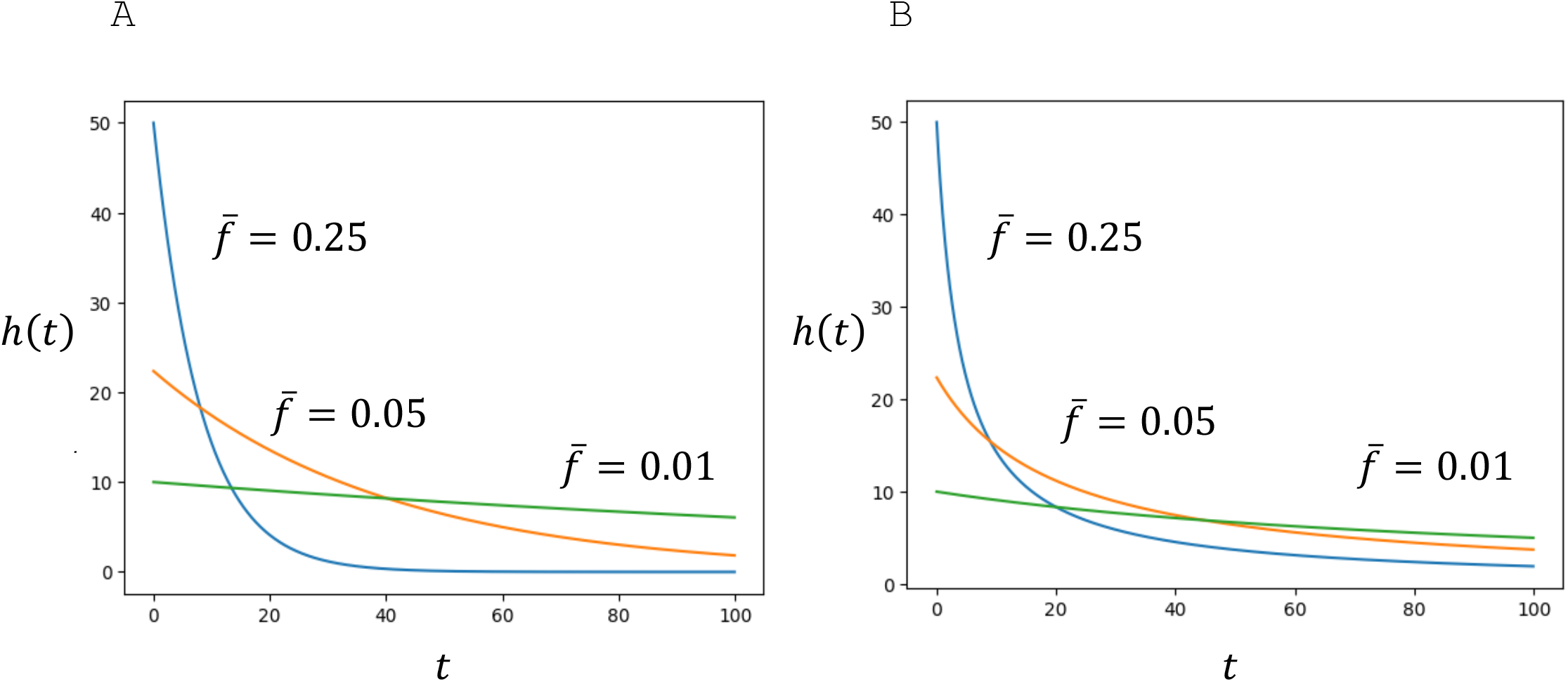
Optimal schedule of naive T cell production *h*(*t*) when all the strains have the same encounter rate 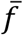. The horizontal axis indicates age *t*. (A) *h*(*t*) when all strains have the same encounter rate 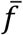. The formula is given by Eq. (8a) in the text. Different curves correspond to the cases with different 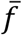, given by Eq. (8a) in the text. A larger 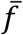 makes the initial level *h*(0) larger, decreases more rapidly, and the total production of naive T cells (area under the curve) larger. (B) *h*(*t*) when the encounter rate *f* has a large variance between strains, given by an exponential distribution (*a* = 1). *h*(*t*) decreases with age *t*, following Eq. (8b), which is a power function. Other parameters: *Kn*/*m* = 10^4^.

The initial rate of naive T cells is 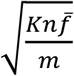, which is high when encountering new pathogen is risky (large *K*), cost of maintaining the thymus is small (small *m*), and the total rate of encounter with pathogen strains in the environment is high (large 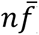). *n* is the diversity of pathogen strains and 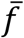 is the rate of encountering with each pathogen strain. The optimal rate of naive T cell production for infants is high in the environment with many kinds of pathogens, each being abundant.

The rate of decrease in *h*(*t*) with age is fast if the rate of encountering with pathogens is high (large 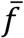). This is because the host acquires immune memory quickly in the environment where each pathogen is abundant. The formula Eq. (3) predicts that thymic involution occurs rapidly in unsanitary environments and more slowly in hygienic environments.

The total amount of naive T cell production is expressed as follows:

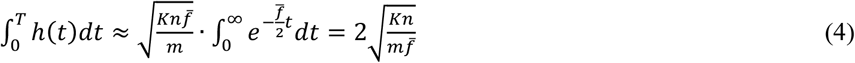

where we approximate the maximum age by infinity, which is an acceptable approximation because the rate of supply in naive T cells decline quickly with age. Eq. (4) indicates that the total lifetime number of naive T cells formed increases with the risk of fighting with a novel pathogen infection and the diversity of pathogens, while decreasing with the cost of maintaining thymic activity and the rate of encountering each strain of risky pathogens.

These results suggest that, if we compare areas with the same diversity of strains (*n* is the same), in the environment full of pathogenic antigens (i.e., not very hygienic environment), as the rate of encountering with the same strain 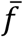 is larger, the optimal rate of naive T cell production *h*(*t*) starts higher, but it decreases more quickly with age; and the total number of naive T cells to produce is smaller.

### 3.2 Ratio of fighting cost to maintenance cost

We note that the optimal rate of naive T cell production is enhanced when the risk of the fighting *K* is high and the cost of maintaining the ability to produce naive T cells *m* is small. When their ratio 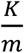 increases, the number of naive T cell production increases for all ages. Hence, the total number of naive T cell to produce in lifetime also increases in proportion to the square-root of the ratio 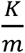.

### 3.3 Diversity of pathogen strains

The dependence of *h*(*t*) on *n*, the diversity of pathogen strains or the total kinds of pathogen strains, is the same as its dependence on 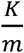. Hence, we conclude that the optimal rate of naive T cell production is higher when the diversity of pathogen strains is greater, irrespective of the host’s age.

## 4. When antigens differ in their frequency of encounter

Quantity *f*_*i*_ indicates the rate of encountering with the antigen that requires host’s adaptive immunity operation. This may differ between antigens. The sum given by Eq. (2) is a mixture of exponential functions with different rates of decrease. In this section, we consider the effect of differences in *f*_*i*_ among pathogen strains.

If we plot *logh*(*t*) over age *t*, it is represented by a curve with a negative slope and the rate of decrease is faster for young ages and becomes slower for older ages. This is proved mathematically because 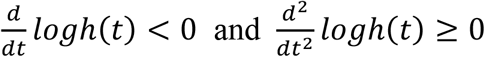 hold. Refer to Appendix A for the derivation.

### 4.1 Gamma distribution

We consider the case that the distribution of *f*_*i*_ can be approximately by a continuous probability distribution of *f*, denoted by *ψ*(*f*). Suppose that *ψ*(*f*) is close to a gamma distribution 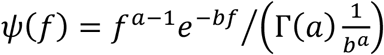, which is a probability distribution over positive value (*f* > 0) with a single peak (if *a* > 1) or the one with a decreasing probability density (if 0 < *a* ≤ 1). It has mean 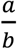 and variance 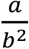. Then, we have the following approximation:

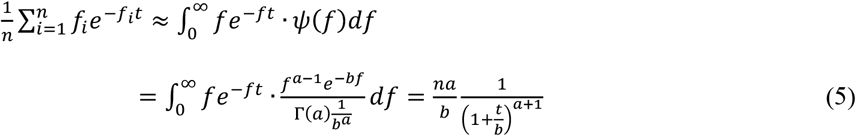

Refer to Appendix A for the calculation. Hence, the optimal rate of naive T cell production, given by Eq. (2), becomes as follows:

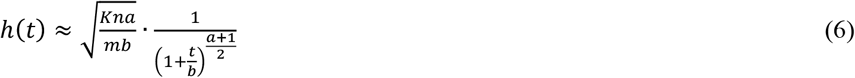

This indicates that the optimal size of the naive T cell supply rate decreases with age, not exponentially, but according to a power function.

By noting that the mean value of *f* is 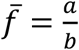, we can rewrite Eq. (6) as follows:

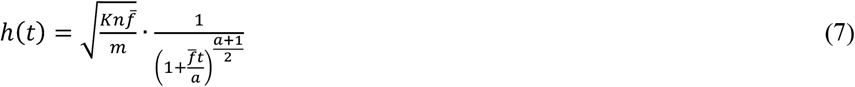

where we removed parameter *b* by introducing the mean 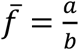. The ratio of variance to the squared mean, given by *variance*/*mean*^2^ = 1/*a*, is independent of the choice of unit for *f*. The ratio 1/*a* provides a good measure for the relative magnitude of variation of the rate *f*. The optimal rate of naive T cell supply *h*(*t*) becomes simplified forms in two different choices of *a*, as shown below:

#### (i) When *a* is very large

In the limit of *a* → ∞, we have the following result:

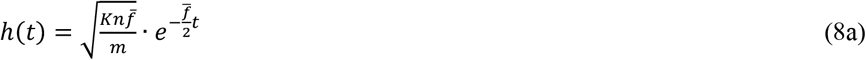

because 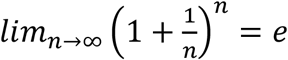. Eq. (8a) is the same as Eq. (3) when the values of *f*_*i*_ between different antigens are the same.

#### (ii) When *a* is equal to unity

By setting *a* = 1, the gamma distribution becomes an exponential distribution with mean 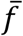 (i.e. 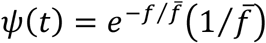). Then we have the following simple formula:

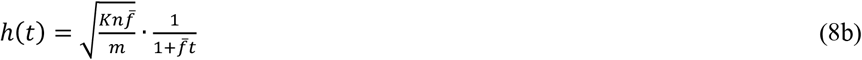

which decreases as a power function (or a hyperbolic function) of age *t*. This corresponds to the case that the value of *f*_*i*_ of different antigens is close to an exponential distribution with mean 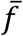 but a sufficiently large variance. An example is illustrated in Fig. 2B.

If 1 < *a* < ∞, we have an intermediate situation between exponential and power functions. Please refer to Appendix A for details.

In Fig. 3, three columns, A, B, and C, illustrate the cases with *a* = 100, *a* = 10, and *a* = 1, respectively. Top parts indicate distribution of *f*. Fig. 3A, 3B, and 3C have the same mean 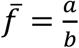 but different variance 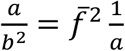. The variance is small for Fig. 3A, intermediate for Fig. 3B, and large for Fig. 3C. Note that in Fig. 3C the probability distribution of *f* is an exponential distribution. Bottom parts indicate *logh*(*t*), the optimal supply rates in logarithmic scale. We can see that the slope of *logh*(*t*) is negative for all ages, implying that *h*(*t*) decreases with age. The rate of decrease in *logh*(*t*) is faster for young ages and becomes slower for older ages. Hence the curve of *logh*(*t*) has a positive second derivative as a function of *t*, which can be shown mathematically (see Appendix A). The straight lines connect two points at age 0 and age 80 and are given as follows:

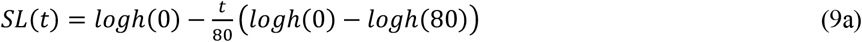

**Figure 3.**
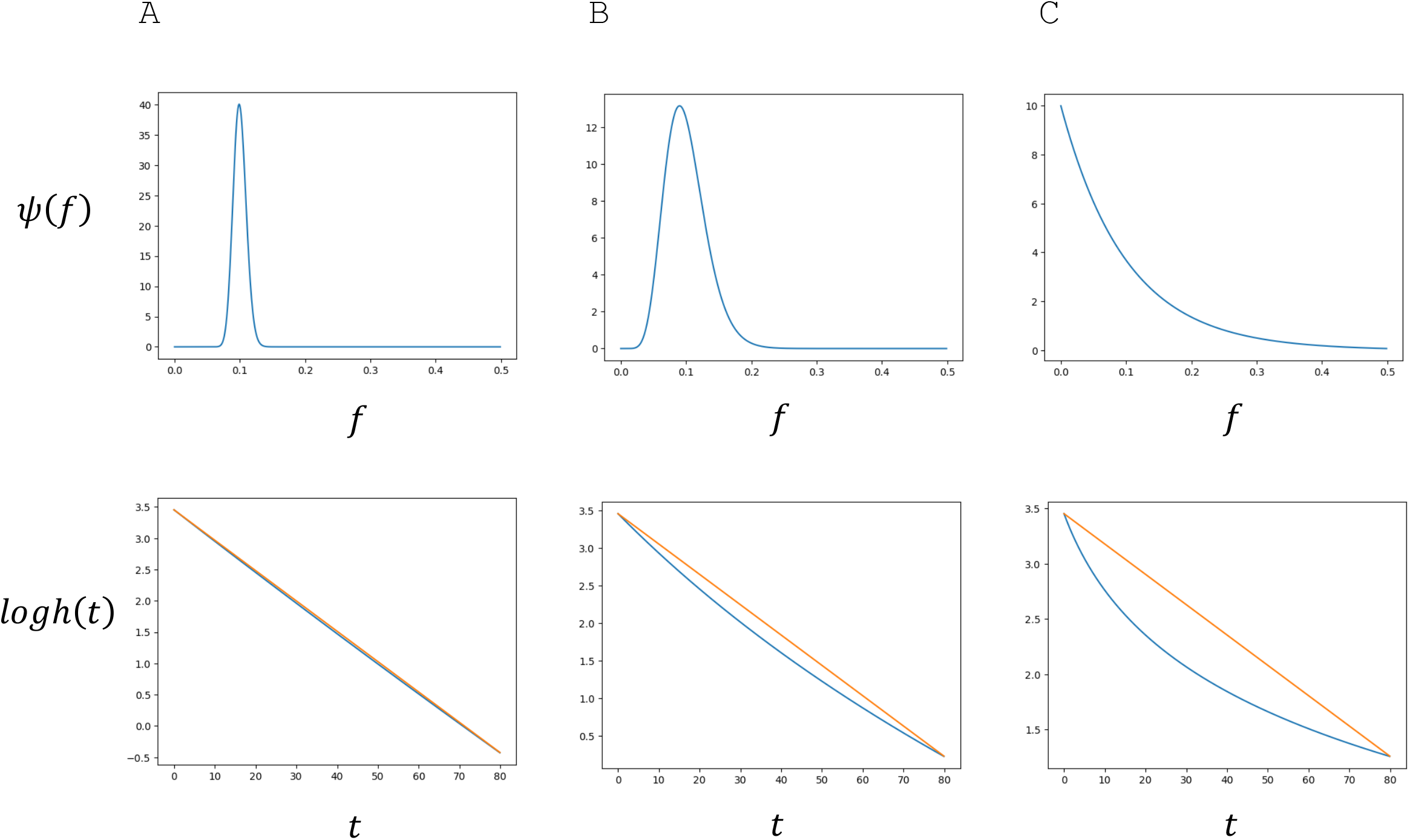
Optimal schedule of naive T cell production *h*(*t*) for different probability distribution of *f*. The encounter rate *f* of different strains follows Gamma-distributions with probability density *ψ*(*f*) = *g*_*a,b*_(*f*). Three cases, (A), (B), and (C), have the same mean value 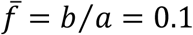. They however differ in variance of *f*. Ratio, *variance*/*mean*^2^ = 1/*a*, is as follows: (A) 0.01; (B) 0.1; (C) 1. The parts in the left column represent different shapes of the probability distribution. The parts in the middle column show *logh*(*t*). The horizontal axis indicates the age. The parts in the right column indicate 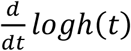, which is a nearly constant in (A), but increases with age *t* in (B) and (C). Parameters are: *Kn*/*m* = 10^4^. (A) *a* = 100, *b* = 1000. (B) *a* = 10, *b* = 100. (C) *a* = 1, *b* = 10.

We consider the maximum difference between the straight line and the curve as follows: *max*_0<*t*<80_[*SL*(*t*) − *logh*(*t*)]. We considered the relative magnitude of the maximum difference divided by (*logh*(0) − *logh*(80)). It is defined as follows:

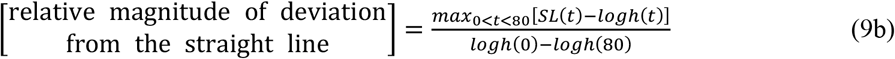

In Fig. 4. we plot the relative magnitude of deviation, given as Eq. (9b). The horizontal axis is 1/*a*, ratio of variance to squared mean: *var*/*mean*^2^. Fig. 4B illustrates that as the variance increases, the curve of *logh*(*t*) deviates more from the straight line, and the magnitude of difference increases with 1/*a*. Refer to Appendix A for details.

**Figure 4.**
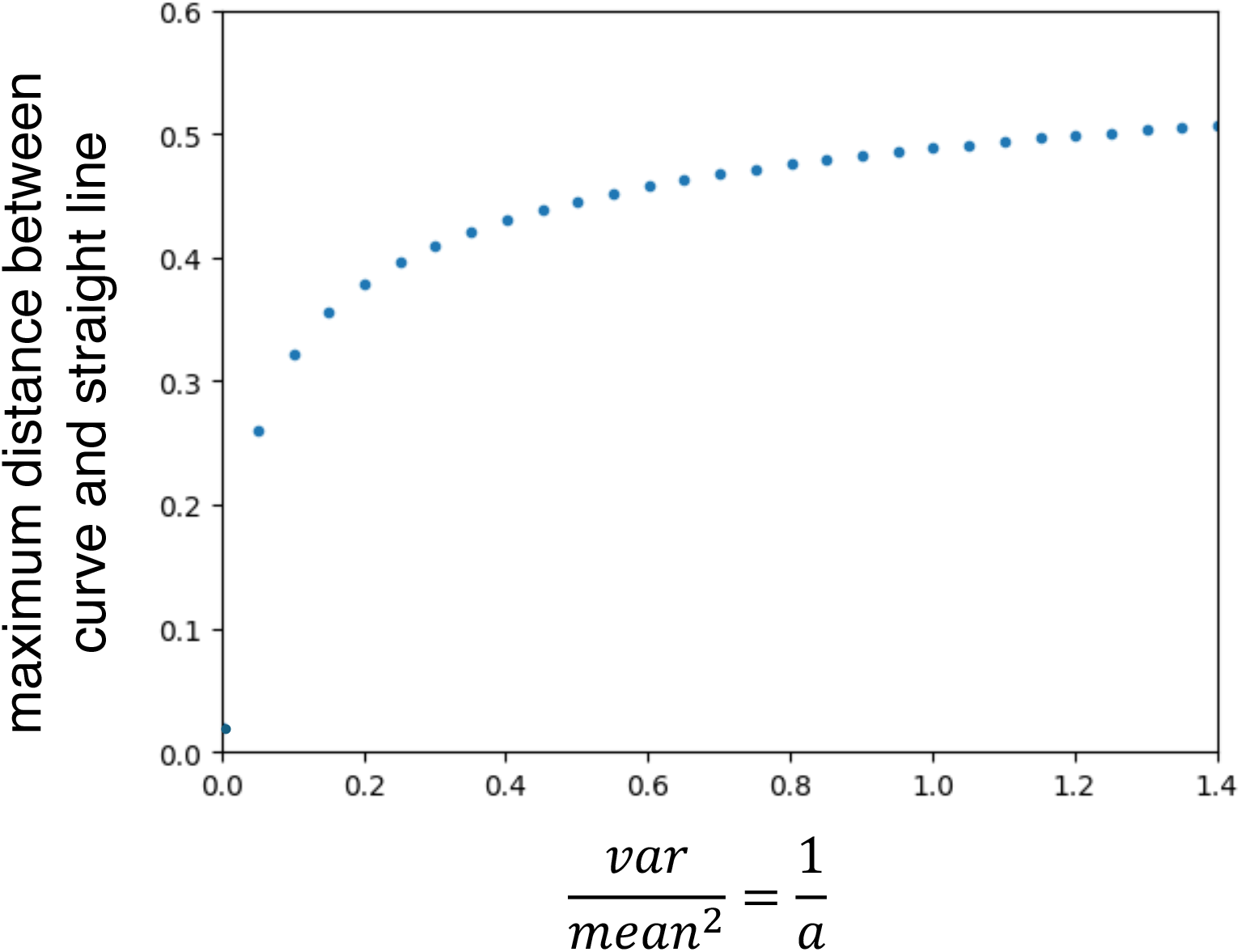
The magnitude of the deviation of logarithmic *h*(*t*) from exponential decrease. The encounter rate *f* follows a gamma distribution. The vertical axis is the relative magnitude of deviation, defined as Eq. (9b). Relative magnitude of the different between the curve and the straight line. The horizontal axis represents *variance*/*mean*^2^ = 1/*a*. The parameter *f* among antigens is given by a gamma distribution. Mean is fixed: 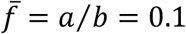. *Kn*/*m* = 10^4^.

### 4.2 Lognormal distribution for antigen-abundance relation

In ecology, a “community” refers to an assembly of different species that inhabit a specific area and coexist while interacting with one another. A commonly observed species-abundance relation in a well-studied community is a lognormal distribution (Preston 1948; Pielou 1966; Whittaker 1975). It states that the fraction of species with the logarithmic abundance *z* = *logf* has a normal distribution with mean *μ* and variance *σ*^2^. The probability distribution of *f* = *e*^*z*^, instead of *logf* = *z*, is given as follows:

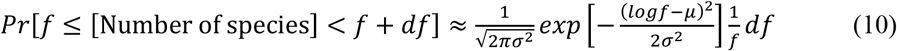

This is *ψ*(*f*), the probability distribution of *f* between strains. The mean value of *f* is 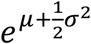 and the variance is 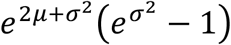.

From Eq. (2), we have

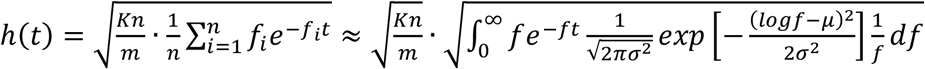

The integral within the square root is rewritten as follows:

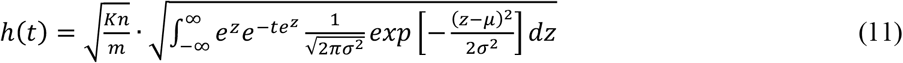

where we set *z* = *logf*.

Fig. 5 illustrates the optimal speed of naive T cell supply from the thymus *h*(*t*), shown in Eq. (11). Fig. 5A, 5B, and 5C illustrate the probability distribution of *f* (top parts) and the logarithmic value of *h*(*t*), namely *logh*(*t*) (bottom parts). They have the same mean values 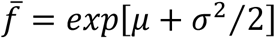, but different variances. The naive T cell supply *h*(*t*) decreases with age *t*. The bottom figures of Figs. 5A, 5B, and 5C illustrate *logh*(*t*) decreases with age with a curve. When *σ*^2^ is small (Fig. 5A), the distribution of *f* is sharpy centered around 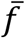, and the optimal *h*(*t*) is close to an exponential function, indicated by a straight line. As *σ*^2^ becomes larger (Figs. 5B, 5C), the distribution of *f* has a larger variance, and *h*(*t*) becomes more like a power function of age. The curve of *logh*(*t*) is deviated from a straight line more strongly as variance *σ*^2^ increases.

**Figure 5.**
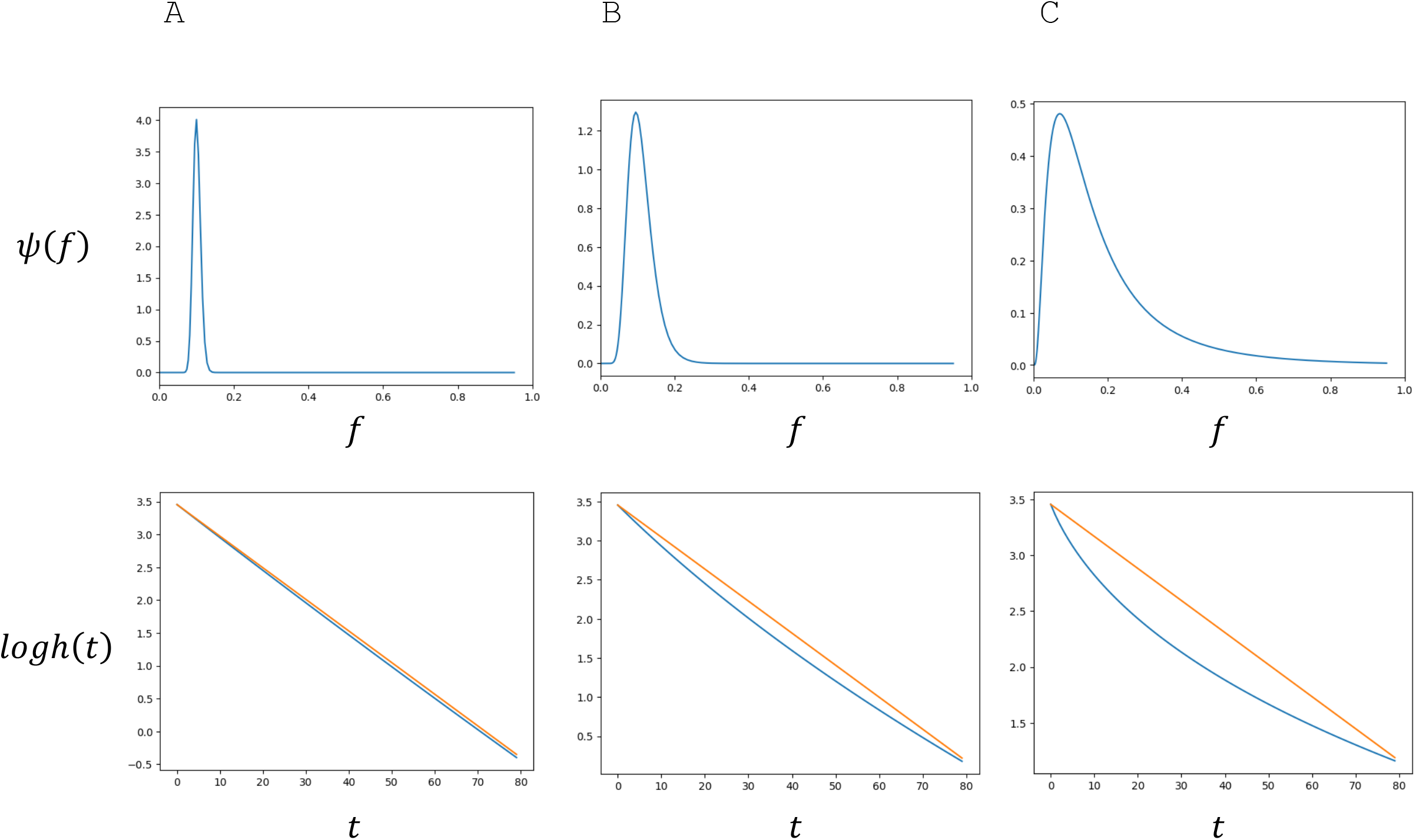
Optimal schedule of naive T cell production *h*(*t*) for different probability distribution of *f. ψ*(*f*) follow log-normal distributions. Three cases, (A), (B), and (C), have the same mean value 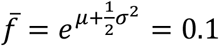; but differ in variance. Ratio, 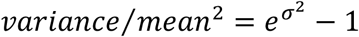, is as follows: (A) 0.01; (B) 0.1; (C) 1. The parts in different columns are the same as in Fig. 3. Parameters are: *Kn*/*m* = 10^4^. (A) *μ* = −2.30495, *σ*^2^ = 0.0099. (B) *μ* = −2.3475, *σ*^2^ = 0.0995. (C) *μ* = −2.645, *σ*^2^ =0.69.

Fig. 6 indicates relative magnitude of the difference between the curve and the corresponding straight line, given by Eq. (9b). The line connects (0, *logh*(0)) and (80, *logh*(80)), and the maximum difference between *logh*(*t*) and *SL*(*t*) is divided by the difference, *logh*(0) − *logh*(80). The relative magnitude of the difference increased with ratio *variance*/*mean*^2^ of *f*.

**Figure 6.**
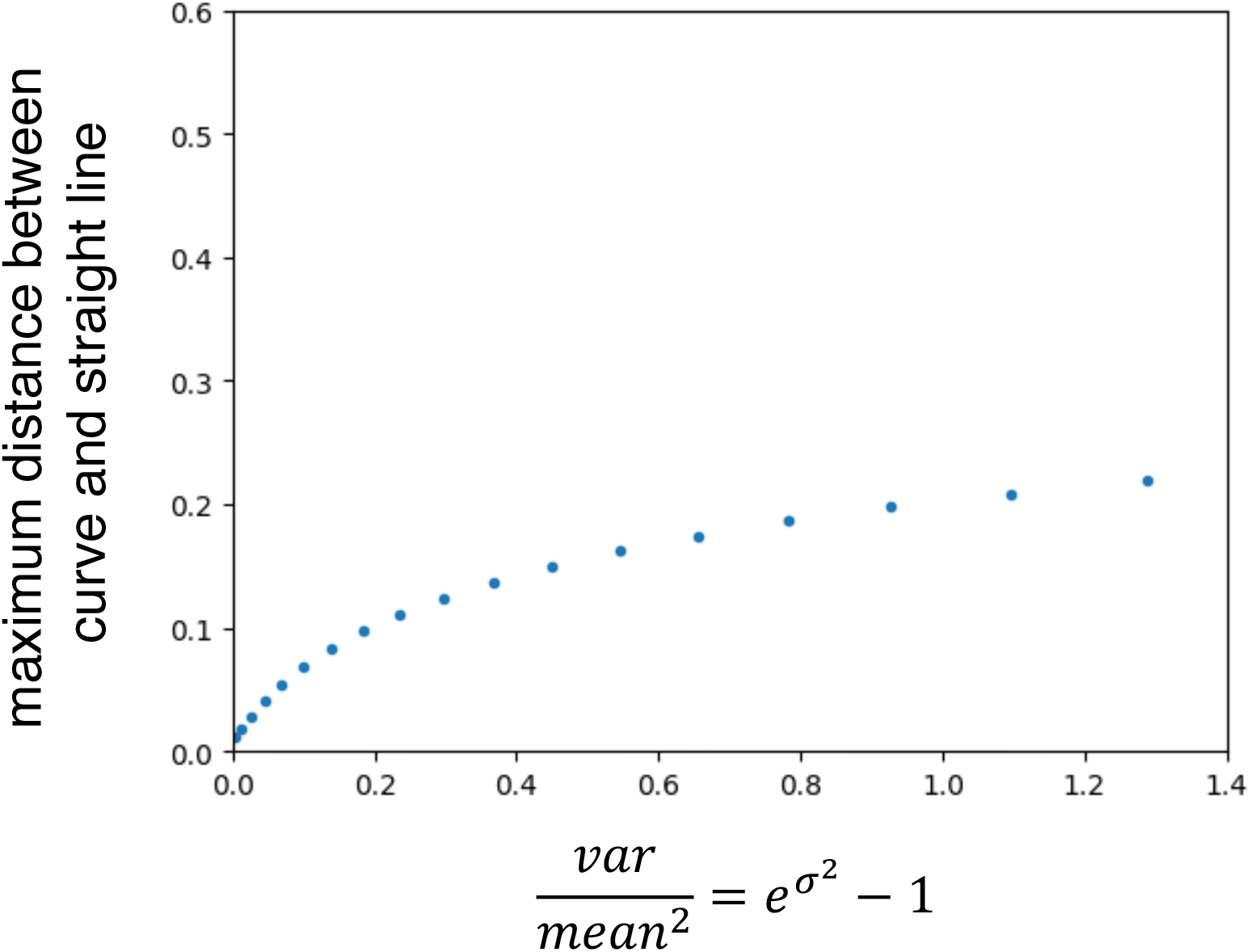
Relative magnitude of the different between the curve and the straight line. It is defined by Eqs. (9a) and (9b). The horizontal axis represents 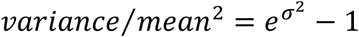. The parameter *f* among antigens is given by a log-normal distribution. Both *μ* and *σ*^2^ changed with the mean fixed: 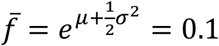. *Kn*/*m* = 10^4^

## 5. Discussion

In this study, we examine thymic involution, represented by the age-dependent decrease in the rate of naive T cell supply, as an adaptive life history schedule. The reduction in the ability to produce naive T cells is the optimal strategy in response to the age-dependent decrease in the need for the capability to cope with novel antigens, which results naturally from the increase of immune memory due to exposure to infectious events.

Most literatures on thymic involution regard it as an example of degradation of biological function with age in the immune system, and many refer to it as “immuno-senescence” (Sidler et al. 2013; Goronzy et al. 2015). The term *senescence* would be suitable for describing the degradation of immune function as an individual passes reproductive age, as aging is a progressive decline in organ or tissue function caused by the accumulation of cellular damage (Sun et al. 2023). However, it does not apply to changes occurring from infancy to adolescence, which take place before reproductive ages.

In this study, we examine thymic involution as an example of adaptive growth schedule considering the cost of maintaining the ability to cope with novel antigens that might potentially be pathogenic. We formalized the problem of the optimal life history schedule. We constructed the simplest model for the cost of fighting with novel antigens, which are potential pathogens, and the cost of maintaining thymic function to produce naive T cells, which are important for the adaptive immunity in learning how to suppress these novel antigens. The results are as follows:

First, when the pathogen strains are similar in encounter rate, the optimal schedule of naive T cells should decline as an exponential function of age. There are literatures supporting exponential decline of the number of thymic cells with age (Steinmann et al. 1985; Bar-Dayan et al. 1999; Kulesh et al. 2024), which release naive T cells decreasing with age. Although cell proliferation may increase the total number of naive T cells in the periphery, it does not enhance the diversity of antigen specificity among them. The ability to detect novel antigens (pathogens) depends on the diversity of naive T cells, which is regulated by the number of newly formed naive T cells produced by the thymus. Some literature concluded the exponential decline of the naive T cell output from the thymus starts early in life (Douek et al. 1998; Kulesh et al. 2024), although others stated the rate of decline slows down at later ages (Ritter and Palmer 1999; Steinmann 1986; Mitchell et al. 2010).

Second, when the rate of pathogen encounters in the environment is higher, the optimal production rate of naive T cells at birth increases; however, this rate declines more rapidly with age. As a result, the total number of naive T cells producing in lifetime would be smaller. It would be worthy to search for literature supporting or refuting this prediction. The thymic function of producing naive T cells continues even in middle- and elder ages (Bar-Dayan et al. 1999).

Third, if the risk of fighting with a novel pathogen by the adaptive immunity is greater and if the cost of maintaining the thymic function of producing naive T cell is smaller, the number of naive T cell production increases at all ages. The same prediction holds for the case of a higher diversity of pathogens in the environment.

Fourth, if pathogens are clearly different with each other concerning the rate of encounter with the host, the optimal rate of naive T cell production decreases as a power function, rather than an exponential function, of age. The exponential rate of decrease is rapid for young ages but slower in older ages. There is some literature concluding that the rate of decline in naive T cell supply is faster at young ages and slows down as individuals get older (Ritter and Palmer 1999; Steinmann 1986; Mitchell et al. 2010), suggesting a power function of age.

### 5.1 Study of alternative scenarios

In this study, we analyzed the simplest possible scenario, but future studies should examine alternative scenario for the thymic involution or extend the model to address a broader range of assumptions. For instance, in this paper, we assumed that the decline in thymic function is attributed to the reduced rate of encounters with novel antigens as the host ages. We examined the scenarios where adaptive immunity must be activated to combat pathogens, leading to the formation of immune memory and the elimination of the pathogen. An important advantage of adaptive immunity in combating pathogen infections is the formation of immune memory, which allows the host to respond more effectively to reinfections by the same pathogen. However, if the likelihood of re-encountering the same antigens is low, relying on innate immunity to suppress pathogens might be more advantageous than activating the adaptive immune response. This suggests that adaptive immunity may be more likely to be advantageous for young individuals than for older ones, providing an alternative rationale for why thymic activity decreases with age. It would be worthwhile to examine the conditions under which this reasoning holds true in future studies.

We assumed that the cost of maintenance is proportional to the rate of naive T cell production, multiplied by a constant *m*. However, the same level of investment in the thymus might represent a much heavier cost to an infant with a small body size compared to an adolescent or an adult with a larger body size. According to life history strategy theory, the cost of investing in defense activities should be measured as the marginal impact on future reproductive success or fitness (Trivers 1972). This perspective highlights the strong age dependence of the cost of such activities. If body size increases very rapidly, an investment in defense activity equivalent to one gram of carbon by a small individual may correspond to an investment of 100 grams of carbon by a larger individual.

Naive T cell receptor repertoires exhibit a wide distribution of clone sizes (de Greef et al. 2020). We considered multiple strains that may differ in their overall abundance, but we assumed that these abundances remain constant over time. In reality, strains that are highly abundant in the early stages of infection may become less abundant later. Investigating the temporal dynamics of pathogenic strains represents an important direction for future theoretical research.

The current study explores the general trend of age-related declines in the interest and capability in learning novel and unfamiliar experiences. In general, responding to new phenomena requires navigating various challenges before we learn effective methods to cope with them. Maintaining the ability to address novel threats comes with a cost, as it necessitates sustaining interest in new topics. As we age and learn to respond based on past experiences, we become better at avoiding the costs associated with new experiences, thus reducing the costs of maintaining the ability to acquire further knowledge. In personality psychology, openness refers to an individual’s willingness to consider new ideas, experiences, and possibilities (Roberts et al. 2006). Individuals high in openness demonstrate a strong desire to learn, a readiness to try new things, and enthusiasm for creative activities and novel ideas. Openness reflects a person’s ability to adapt to new challenges as they arise. Younger individuals typically exhibit higher levels of openness, but as they age, they may become more set in their ways, leading to a decline in openness (Roberts et al. 2006). This age-related decrease in openness might be understood as an adaptive strategy, comparable to thymic involution examined in this study.

## Appendix A

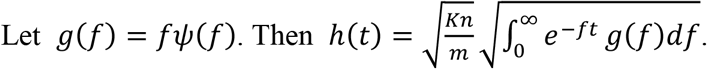

Using Eq. (2), we have the following:

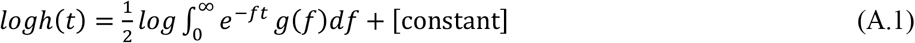

By calculating the derivative with respect to *t*,

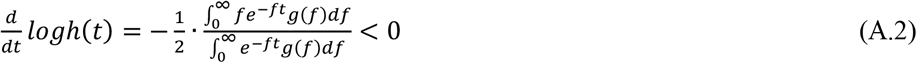

which indicates that logarithmic value of *h*(*t*) has a negative slope and it decreases with age.

The derivative of Eq. (A.2) becomes

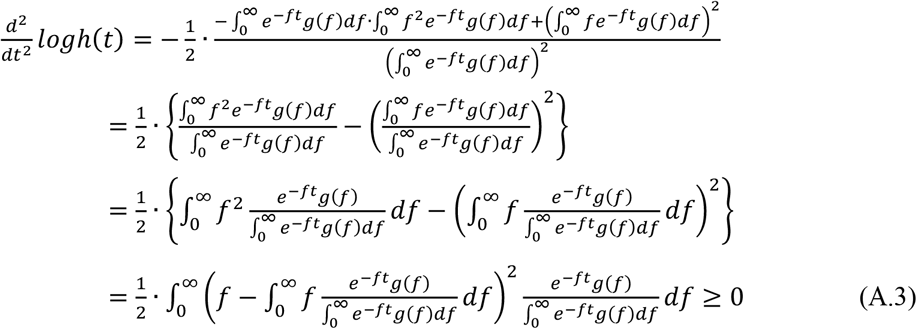

This implies that the slope of the graph increases with age *t*. The speed of decrease is fast initially and becomes slower as ages.

### 4.1 When f_i_ differs between allergens

We here assume that the value of *f* among strains a gamma-distribution

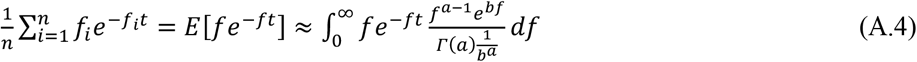

*E*[·] indicates the average with respect to *f*, which follows a gamma distribution 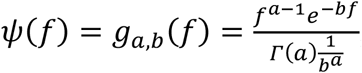, which has the mean and variance as follows: 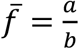 and 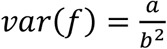

The ration of variance to squared mean is

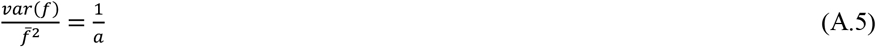

which indicates the relative magnitude of variance. Note that it does not change even if we change “unit” of *f*.

The integral Eq. (A.1) is rewritten as follows:

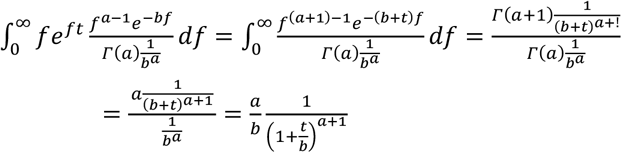

The optimal supply rate of naive T cells from the thymus Eq. (2) becomes as follows:

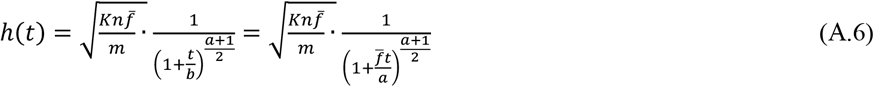

We consider the cases with the mean fixed as 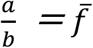, but ratio of variance to squared mean 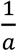 is fixed. When *a* is very large, the variance is very small, and the distribution of *f* has a sharp peak around 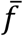. In contrast, when *a* is small, the distribution of *f* has some variance. In the following, we calculate the dependence on the value of *a*.

[Case I] When *a* becomes infinitely large, since

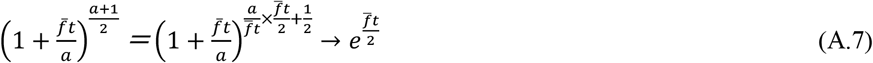

*h*(*t*) becomes as follows:

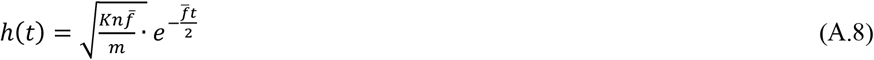

which decreases with age *t* as an exponential function.

[Case II] When *a* = 1, the probability density of *f* is an exponential distribution 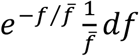. Then, we have the following result for the optimal rate of naive T cell output from the thymus:

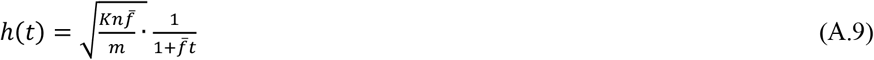

which decreases with age *t* as a power function.

[Case III] If 1 < *a* < ∞, we have an intermediate situation between exponential and power functions.

We adopted these formulas for drawing Figs. 3 and 4.

#### A.2 When f_i_ distributes a log-normal distribution

In community ecology, the species abundance relation that are often observed in the well-studied forest is a log-normal distribution (Preston 1948; Pielou 1966). If we approximate it as a probability density of *f*, the abundance for a randomly chosen species, it is described as follows

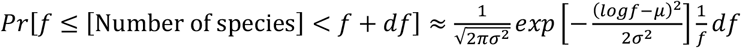

which has mean 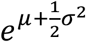 and the variance 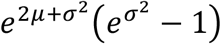. Ration of the variance to squared mean is 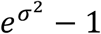, which is an increasing function of *σ*^2^

Then the optimal rate of naive T cell supply from the thymus given by Eq. (2) is written as follows:

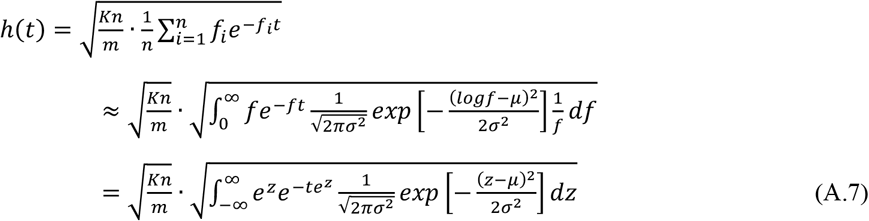

We adopted the last formula for calculating Figs. 5 and 6.

## CRediT authorship contribution statement

Yoh Iwasa: Conceptualization, Methodology, Formal analysis, Visualization, Writing – original draft.

Kosei Matsuo: Conceptualization, Numerical analysis, Visualization, Writing – original draft.

## Acknowledgements

We thank Rena Hayashi, Yuna Kotsubo, Akiko Satake, and Sou Tomimoto for their very helpful comments.

## Data availability statement

The datasets used and/or analyzed during the current study available from the corresponding author on reasonable request.

## References

Appay, V., Sauce, D. (2014) Naive T cells: the crux of cellular immune aging? Experimental Gerontology 54:90–93. doi: 10.1016/j.exger.2014.01.003.

Bains, I., Antia, R., Callard, R.,, Yates, A.J. (2009) Quantifing the development of the peripheral naive CV4+ T cell pool in humans. Blood 113:5480–5487.

Bar-Dayan, Y., Afek, A., Bar-Dayan, Y., Golberg, I., Kopolovic, J. (1999) Proliferation, apoptosis and thymic involution. Tissue and Cell 31: 391–396.

Collins, C.P., Khuat, L.T., Sckisel, G.D., Vick, L.V., Minnar, C.M., Dunai, C., Le, C.T., Curti, B.D., Crittenden, M., Merleev, A., Sheng, M., Chao, N.J., Maverakis, E., Rosario, S.R., Monjazeb, A.M., Blazar, B.R., Longo, D.L., Canter, R.J., Murphy, W.J., (2024) Systemic immunostimulation induces glucocorticoid-mediated thymic involution succeeded by rebound hyperplasia which is impaired in aged recipients. Frontiers in Immunology 15:1429912.

de Greef, P.C., Oakes, T., Gerritsen. B., Ismail, M., Heather, J.M., Hermsen, R., Chain, B., de Boer, RJ. (2020) The naive T-cell receptor repertoire ha an extremely broad distribution of clone sizes. eLife 9:e49900. doi: 10.7554/eLife.49900

Douek DC, McFarland RD, Keiser PH, Gage EA, Massey JM, Haynes, B.F., Polis, M.A., Haase, A.T., Feinberg, M.B., Sullivan#, J.L., Jamieson, B.D., Zack, J.A., Picker, L.J., Koup, R.A. (1998) Changes in thymic function with age and during the treatment of HIV infection. Nature 396: 690–695. 10.1038/25374

Fujiwara, M., Yonezawa, R., Arai, R., Yamamoto, I., Ohtsuka, H., (2012) Alterations with age in peripheral blood lymphocyte subpopulations and cytokine synthesis in beagles. Veterinary Medicine: Research and Reports 3:79–84.

Ganusov, V.V., de Boer, R.J. (2007) De most lymphocytes in humans really reside in the gut? Trends in Immunology 28: 514518.

Goronzy, J.J., Fang, F., Cavanagh, M.M., Qi, Q., Weyand C.M. (2015) Naive T cell maintenance and function in human aging. Journal of immunology 194: 4073–4080.

Hamilton, W.D. (1966) The moulding of senescence by natural selection. Journal of Theoretical Biology 12:12–45. doi: 10.1016/0022-5193(66)90184-6.

Kulesh, V., Peskow, K., Helmlinger, G., Bocharow, G. (2024) An integrative mechanistic model of thymocyte dynamics. Frontiers in Immunology 15:1321309.

Lewkiewicz, S., Chuang, Y.L., Chou, T. (2019) A mathematical model of the effects of aging on naive T cell populations and diversity. Bulletin of Mathematical Biology 81: 2783–2817.

Mitchell, W.A., Long, P.O., Aspinall, R. (2010) Tracing thymic output in older individuals. Clinical and Experimental Immunology 161: 497–503.

Parham, P. (2021) The immune system (5th ed.) W. W. Norton & Company

Palmer, S., Albergante, L., Blackburn, C.C., Newman, T.J. (2018) Thymic involution and rising disease incidence with age. PNAS 115:1883–1888. doi: 10.1073/pnas.1714478115

Pielou, E.C. (1966) The measurement of diversity in different types of biological collections. Journal of Theoretical Biology 13:131–144

Preston, F.W. (1948). The commonness, and rarity, of species. Ecology 29: 254–283.

Rane, S., Hogan, T., Lee, E., Seddon, B., Yates, A.J. (2022) Toward a unified model of naive T cell dynamics across the lifespan. eLife 11:e78168.

Ritter, M.A., Palmer, D.B. (1999). The human thymic microenvironment: new approaches to functional analysis. Seminar Immunol. 111:13–21.

Roberts, B.W., Walton, K.E,. Viechtbauer, W. (2006) Patterns of mean-level change in personality traits across the life course: A meta-analysis of longitudinal studies Psychological Bulletin 132: 1–25. DOI10.1037/0033-2909.132.1.1

Sidler, C., Wóycicki, R., Ilnytskyy, Y., Metz, G., Kowalchuk, I, Kovalchuk, O. (2013) Immunosenescence is associated with altered gene expression and epigenetic regulation in primary and secondary immune organs. Frontiers in Genetics 4:e211. 10.3389/fgene.2013.00211

Steinmann, G.G., Klaus, B., Müller-Hermelink, H.K. (1985) The involution of the ageing human rhymic epithelium is independent of puberty. Scand J Immunol 22: 563–575 10.1111/j.1365-3083.1985.tb01916.x

Steinmann, G.G. (1986) Changes in the human thymus during aging. Curr. Top Pathol. 75:43–68.

Sun, B., Meng, X., Li, Y., Li, Y., Liu, R., Xiao, Z. (2023) Conditioned medium from human cord blood mesenchymal stem cells attenuates age-related immune dysfunctions. Frontiers in Cell and Developmental Biology doi: 10.3389/fcell.2022.1042609

Trivers, R.L. (1972). Parental investment and sexual selection. In B. Campbell (Ed.), Sexual selection and the descent of man, 1871-1971 (pp. 136–179). Chicago, IL: Aldine.

Whittaker, R.H. (1975) Communities and Ecosystems. 2nd Revise Edition, MacMillan Publishing Co., New York.

